# The Human Brain Connectome Weighted by the Myelin Content and Total Intra-Axonal Cross-Sectional Area of White Matter Tracts

**DOI:** 10.1101/2023.03.01.530710

**Authors:** Mark C. Nelson, Jessica Royer, Ilana R. Leppert, Jennifer S.W. Campbell, Simona Schiavi, Hyerang Jin, Shahin Tavakol, Reinder Vos de Wael, Raul Rodriguez-Cruces, G. Bruce Pike, Boris C. Bernhardt, Alessandro Daducci, Bratislav Misic, Christine L. Tardif

## Abstract

A central goal in neuroscience is the development of a comprehensive mapping between structural and functional brain features. Computational models support *in vivo* investigation of the mechanisms mediating this relationship but currently lack the requisite biological detail. Here, we characterize human structural brain networks weighted by multiple white matter microstructural features to assess their potential joint utilization in computational models. We report edge-weight-dependent spatial distributions, variance, small-worldness, rich club, hubs, as well as relationships with function, edge length and myelin. Contrasting networks weighted by the total intra-axonal cross-sectional area and myelin content of white matter tracts, we find opposite relationships with functional connectivity, an edge-length-independent inverse relationship with each other, and the lack of a canonical rich club in myelin-weighted networks. When controlling for edge length, tractometry-derived networks weighted by either tensor-based metrics or neurite density show no relationship with whole-brain functional connectivity. We conclude that structure-function brain models are likely to be improved by the co-utilization of structural networks weighted by total intra-axonal cross-sectional area and myelin content. We anticipate that the proposed microstructure-weighted computational modeling approach will support mechanistic understanding of the structure-function relationship of the human brain.

**AUTHOR SUMMARY:** For computational network models to provide mechanistic links between brain structure and function, they must be informed by networks in which edge weights quantify structural features relevant to brain function. Here, we characterized several weighted structural networks capturing multiscale features of white matter connectivity. We describe these networks in terms of edge weight distribution, variance and network topology, as well as their relationships with each other, edge length and function. Overall, these findings support the joint use of structural networks weighted by the total intra-axonal cross-sectional area and myelin content of white matter tracts in structure-function models. This thorough characterization serves as a benchmark for future investigations of weighted structural brain networks.

## INTRODUCTION

The quest to relate human structural and functional brain networks spans the spectrum of spatial scale and repertoire of data modalities absolutely. At the macroscale, the human brain can be modeled as an anatomical network of discrete neuronal populations (nodes) interconnected by white matter fibers (edges) (Sporns, 2011). Coordinated spatiotemporal patterns of neuronal activity unfolding upon this structural backbone are fine-tuned by white matter microstructure (Hodgkin & Huxley, 1952; Huxley & Stämpfli, 1949; Moore et al., 2020; Pumphrey & Young, 1938) and form the basis of cognition and behavior (Biswal et al., 1995; Greicius et al., 2003; Hampson et al., 2006; Liégeois et al., 2019; S. M. Smith et al., 2009; Martijn P. Van Den Heuvel et al., 2009). Increasingly, MRI facilitates *in vivo* measurement of multi-scale properties of both brain structure (e.g., (Alexander et al., 2019; Drakesmith et al., 2019; Jeurissen et al., 2017; Mancini et al., 2020)) and function (e.g., (Finn et al., 2019; Friston, 2011; Gordon et al., 2017; Liu et al., 2022)). Diffusion MRI streamline tractography and resting-state functional MRI are often respectively used to estimate structural and functional connectivity (SC & FC) networks. Network science provides a framework to bring these fundamentally different substrates into a common space where their features can be quantified (Fornito et al., 2016; Sporns, 2010; Suárez et al., 2020) and used to probe the mechanisms mediating human brain function (e.g., (Cabral et al., 2017; Fornito et al., 2015)).

SC network edges can be weighted by a range of MRI-derived metrics quantifying white matter microstructural features relevant to brain function including voxel-level estimates of tissue diffusivity (e.g., (Caeyenberghs et al., 2016)), neurite density (Zhang et al., 2012), axon diameter distributions (Alexander et al., 2010; Assaf et al., 2008), myelin content (Heath et al., 2018; Mancini et al., 2020), and the g-ratio (ratio of inner/outer diameters of myelinated axons) (Stikov et al., 2011, 2015); as well as tract/bundle-level measures of axonal cross-sectional area (Daducci, Dal Palù, et al., 2015; R. E. Smith et al., 2015). Subsets of these metrics have been investigated using a microstructure-weighted connectomics approach (Boshkovski et al., 2021; Caeyenberghs et al., 2016; Deligianni et al., 2016; Frigo et al., 2020; Mancini et al., 2018; Messaritaki et al., 2021; Schiavi et al., 2020; M. P. van den Heuvel et al., 2010; Martijn P. van den Heuvel & Sporns, 2011; F. C. Yeh et al., 2016), however a comprehensive characterization has not yet been provided.

Our goal is to characterize a range of standard and state-of-the-art weighted structural brain networks in support of their utilization in computational models of brain function. The networks considered here can be grouped into two classes: those computed with tractometry (S Bells et al., 2011) and those computed directly from the streamline weights in a tractogram i.e., streamline-specific. We consider three examples of the latter: (1) the number of streamlines (NoS); and two methods which optimize the streamline weights in a tractogram to increase specificity for white matter structural features (2) spherical-deconvolution informed filtering of tractograms (SIFT2) (R. E. Smith et al., 2015) and (3) convex optimization modeling for microstructure informed tractography (COMMIT) (Daducci et al., 2013; Daducci, Dal Palù, et al., 2015). SIFT2 and COMMIT were designed to overcome known limitations of streamline counts (Girard et al., 2014; Jones, 2010; Jones et al., 2013). While the edge weights in all three networks generally capture white matter features relevant to connection strength, SIFT2 and COMMIT more specifically quantify the total intra-axonal cross-sectional area of white matter tracts (henceforth referred to as “edge caliber”). To date, SIFT2 and COMMIT have not been compared to NoS with uniform connection density (Frigo et al., 2020; Schiavi et al., 2020; C. H. Yeh et al., 2016). Thus, it remains unclear how the edge weights themselves affect network topology.

In contrast, tractometry allows network edge weights to be derived from any volumetric brain image that is co-registered to the tractogram. This increase in methodological flexibility comes at the expense of anatomical specificity. Tractometry is unable to resolve the separate contributions of individual fiber populations to the aggregate value of a voxel. Given that an estimated ∼90% of white matter voxels at typical diffusion MRI resolutions (∼2mm) contain multiple fiber populations (Jeurissen et al., 2012), the quantitative link between white matter microstructure and essentially all tractometry-derived edge weights is biased by partial volume effects.

In this work, tractometry is combined with a diffusion tensor model (Basser, 1995; Basser et al., 1994) to derive networks weighted by FA (fractional anisotropy) and RD (radial diffusivity), which respectively quantify the degree of diffusion anisotropy (i.e., directional dependence) and diffusion magnitude perpendicular to the major axis. The crossing fiber problem described above is also known to limit the ability of diffusion tensor models to quantify white matter features (De Santis et al., 2014; J. D. Tournier et al., 2011). Additional tractometry networks examined here include a network weighted by ICVF (intracellular volume fraction) computed with NODDI (Neurite Orientation Dispersion and Density Imaging) (Zhang et al., 2012), as well as a network weighted by the longitudinal relaxation rate R_1_ (1/T_1_), which has been shown to correlate with histology-derived myelin content (Mottershead et al., 2003).

This characterization of weighted structural brain networks is carried out as follows: (1) within-network features of edge weight distribution and variance; (2) edgewise relationships with FC, edge length and myelin (R_1_); and (3) topological features of small-worldness, rich club and network hubs. Importantly, uniform binary connectivity is enforced across all weighted network variants allowing the edge weights themselves to drive the characterization.

## RESULTS

In 50 healthy adults (27 men; 29.54±5.62 years; 47 right-handed), structural brain networks were estimated from multi-shell diffusion MRI data with probabilistic tractography. Each subject’s structural network was used to compute 8 SC networks in which edges were weighted by: NoS, SIFT2, COMMIT, FA, RD, ICVF, R_1_ and LoS (edge length computed as the mean length of streamlines). The edge weights in NoS, SIFT2 and COMMIT networks were normalized by node volume. Additionally, a static FC network was derived for each subject by zero-lag Pearson cross-correlation of nodewise resting-state time series. Unless otherwise stated, all results shown correspond to networks parcellated with the Schaefer-400 cortical atlas (Schaefer et al., 2018) and include 14 subcortical nodes.

### Structural Brain Networks Vary in the Distribution of Their Edge Weights

Group-level networks weighted by NoS, SIFT2 and COMMIT show spatially distributed patterns of high magnitude edge weights and noticeably accentuate within-module connectivity (**Figure 1A**). Modules correspond to the 7-canonical resting-state networks (Thomas Yeo et al., 2011) plus the subcortex. These patterns are hallmarks of FC networks and are observed in the FC network shown here. The contrast between high and low magnitude edge weights is most evident in COMMIT. By comparison, the spatial variation of edge weight distribution in the tractometry networks is smoother with more pronounced regional concentrations. R_1_ is highest in the edges connecting the visual module to itself and to the rest of the brain; and lowest within the subcortex and between the subcortical and limbic modules. The surface plot shows the highest concentration of R_1_ in the white matter projections of posterior cortical regions.

**Figure 1.**
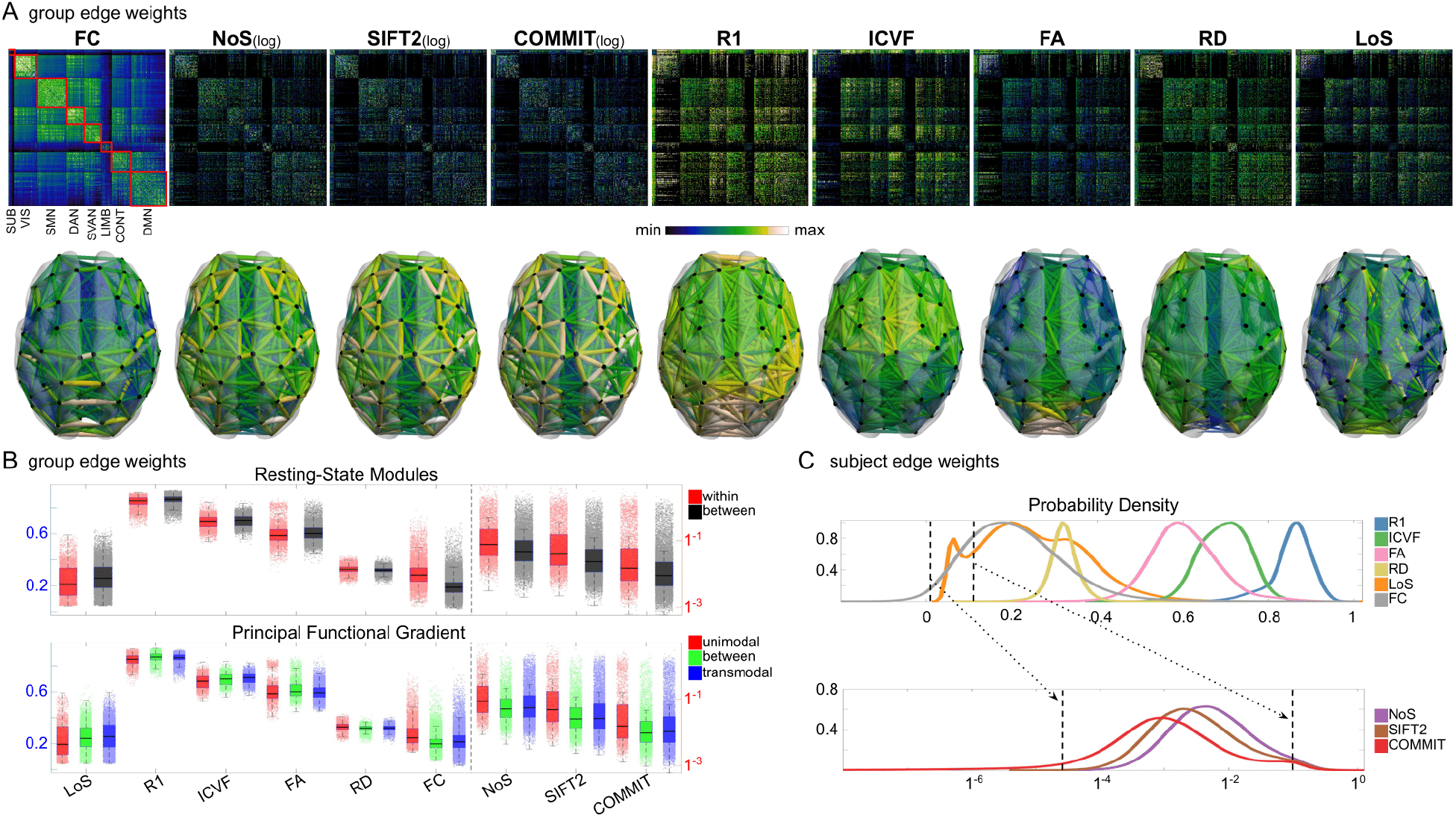
Edge Weight Distribution. (A) Connectivity matrices (top row) of group-level edge weights for FC (functional connectivity), NoS (number of streamlines), SIFT2 (spherical-deconvolution informed filtering of tractograms), COMMIT (convex optimization modeling for microstructure informed tractography), R_1_ (longitudinal relaxation rate), ICVF (intra-cellular volume fraction), FA (fractional anisotropy), RD (radial diffusivity) and LoS (mean length of streamlines). Each network is composed of 414 nodes as defined by the Schaefer-400 cortical parcellation and 14 subcortical ROIs. Nodes are grouped into the canonical resting state modules (Thomas Yeo et al., 2011) plus the subcortex: SUB (subcortex), VIS (visual), SMN (somatomotor), DAN (dorsal attention), SVAN (salience ventral attention), LIMB (limbic), CONT (control), and DMN (default mode). 3D cortical surfaces (bottom row) of group-level edge weights in the Schaefer-100 parcellation generated with BrainNet Viewer (Xia et al., 2013). Edge diameter and color indicate weight magnitude. (B) Distribution of group-level edge weights binned by: (top) within and between module; (bottom) unimodal, transmodal and between. Unimodal is defined as the VIS and SMN modules. Transmodal is defined as the DMN, CONT, DAN and SVAN modules. (C) Probability density of pooled subject-level edge weight distributions. R_1_, ICVF, FA, RD, LoS and FC are shown on a linear x-axis (top), and NoS, SIFT2 and COMMIT are shown on a logarithmic x-axis (bottom). All networks were normalized to the range [0 1] by dividing by the subject-level max for visualization. The edge weights in NoS, SIFT2 and COMMIT networks were log_10_ transformed for these plots.

Group-level edge weight distributions are summarized with respect to two important organizational patterns of brain function (**Figure 1B**): within and between resting state modules (Thomas Yeo et al., 2011); and along the principal functional gradient (Margulies et al., 2016). NoS, SIFT2 and COMMIT mirror FC in both plots with greater edge weight magnitude within module, especially within unimodal modules. R_1_, ICVF, FA and RD generally mirror LoS with the reverse trend: higher between module and lowest in unimodal modules. This suggests that tractometry-derived networks may be influenced by edge length to a greater extent.

Subject-level edge weight distributions in R_1_, ICVF, FA and RD are near-normal and network-specific (**Figure 1C**). They differ in both the magnitude (R_1_ > ICVF > FA > RD) and dynamic range (FA & ICVF > R_1 &_ RD) of their edge weights. In contrast, NoS, SIFT2 and COMMIT distributions are highly skewed and tend to be much lower in magnitude (dashed lines). This effect is greatest in COMMIT suggesting that the optimization performed by COMMIT exerts a stronger scaling effect than SIFT2. These results support the conclusion that the structural networks considered here quantify subsets of white matter features which are at least partially non-overlapping.

### Edge Weights in Streamline-Specific Networks Are More Variable

Edge weight variance was quantified using the Quartile Coefficient of Dispersion (CQD) due to its robustness to outliers and skewed data. The CQD is computed from the 1^st^ and 3^rd^ quartiles as:

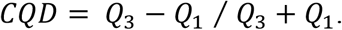

*Intra-subject* variance is roughly 2-fold greater in NoS, SIFT2 and COMMIT than LoS and FC; and an order of magnitude greater than R_1_, ICVF, FA and RD in all subjects (**Figure 2A**). COMMIT is the highest overall. Subjects are more tightly clustered in all weighted SC networks, relative to FC: *intra-subject* CQD values span roughly a 4-fold greater range in FC. This suggests that individual diversity of functional connectivity is not necessarily reflected in the variability of their structural networks. These patterns are repeated for *inter-subject* variance. However, FC shows a small subset of highly variable edges with roughly 4-fold greater CQD than the maximum values observed in COMMIT i.e., the most subject-specific connections are functional. The very low edge weight variability in R1, ICVF, FA and RD is in part due to the widespread smoothing effect (partial voluming) resulting from the tractometry computation.

**Figure 2.**
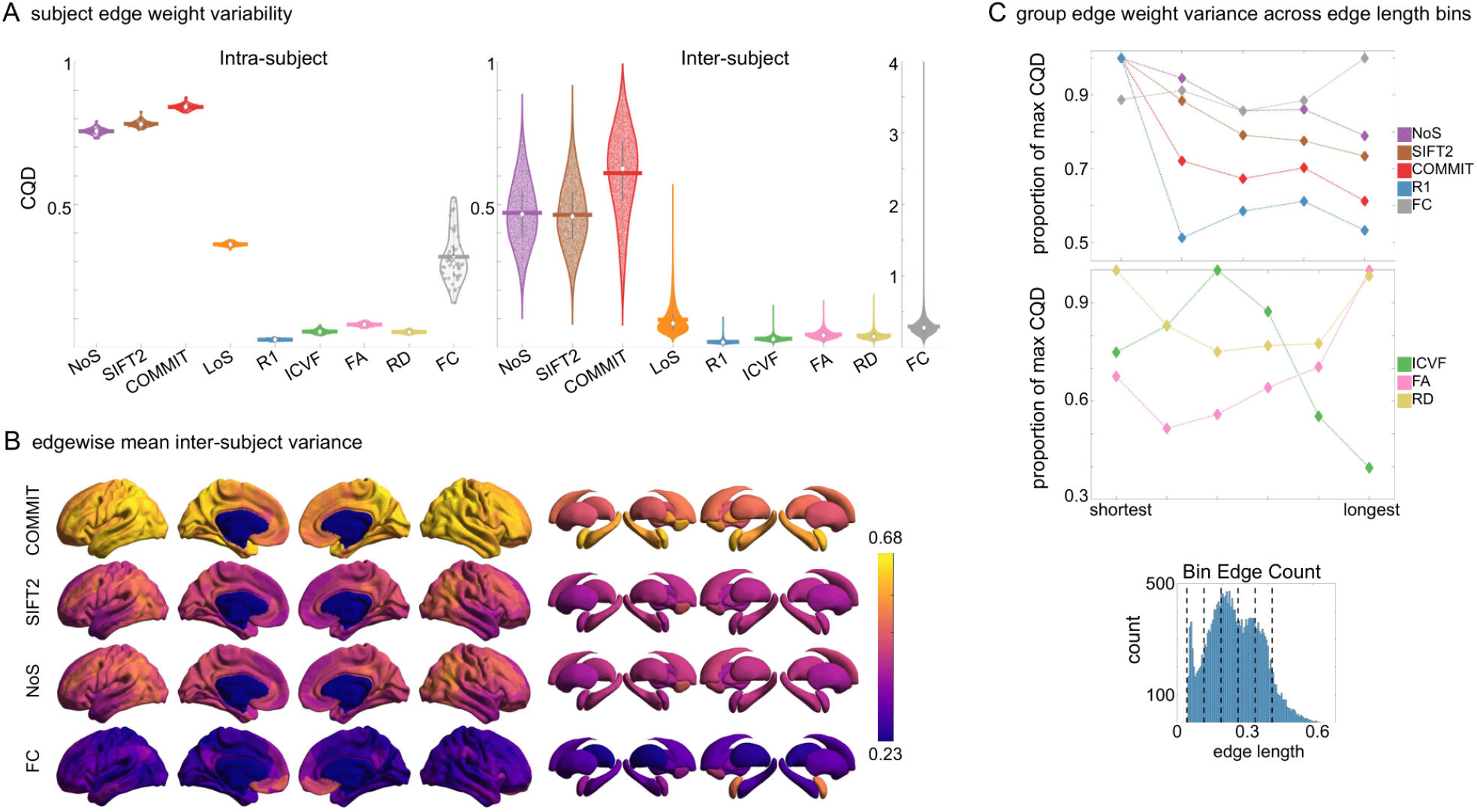
Edge Weight Variability. Variability is quantified using the coefficient of quartile dispersion (CQD). (A) Violin distributions of **intra-subject** (left) and **inter-subject** (right) edge weight variance. Colored data points respectively correspond to individual subjects (N=50) and edges (N=8549). (B) Surface projections of edgewise mean inter-subject variance for cortical nodes in the Schaefer-400 parcellation (left) and 14 subcortical nodes (right). Cortical and subcortical surfaces were respectively generated with BrainSpace (Vos de Wael et al., 2020) and ENIGMA toolboxes (Larivière et al., 2021). (C) The proportion of within-network max CQD is shown across edge length bins for FC, NoS, SIFT2, COMMIT and R_1_ (top), as well as ICVF, FA and RD (middle). Edge weights are grouped into 6 bins according to edge length, as illustrated by the histogram (bottom). The edges of bins 1-5 were linearly spaced of width, w. The edges of the final bin were of width 3w.

In general, *inter-subject* edge weight variance is more spatially distributed in SC networks relative to FC (**Figure 2B**). COMMIT shows the highest mean CQD over the entire cortex and subcortex. NoS, SIFT2 and COMMIT all show lateral-medial and posterior-anterior cortical gradients. Mean CQD in FC shows the highest concentration in medial inferior frontal cortex and to a lesser extent, the expected pattern of high variance in association cortex. The most variable subcortical regions include the hippocampus, amygdala and accumbens.

Many features of brain networks (e.g., connection probability, weight magnitude) are known to vary with edge length. Here, we examined the relationship between edge weight variability and edge length by computing the CQD within subsets of group-level edge weights binned according to their edge length (**Figure 2C**). Edge weight variance in NoS, SIFT2, COMMIT and R_1_ is highest in the shortest edges and decreases with edge length. ICVF roughly follows the same pattern. FA and RD instead show the highest variability in the longest edges. Overall, the edge weights in streamline-specific SC networks (NoS, SIFT2 and COMMIT) show greater contrast both within and across subjects. SC networks show network-dependent relationships between edge weight variance and edge length. Shorter edges are more variable in myelin- and connection strength-weighted networks, and longer edges are more variable in networks with edge weights derived from a diffusion tensor model.

### Opposing Correlations with Function in Connection-Strength-& Myelin-Weighted Networks

Shifting to inter-network edge weight relationships shows that SC networks are differentially related to FC (**Figure 3A)**. Importantly, we also see that all brain networks (SC and FC) are strongly and differentially related to edge length at the subject and group levels. Correlations with edge length are negative for NoS, SIFT2, COMMIT, RD and FC; and positive for R_1_, ICVF, and FA. Correlation magnitude is strongest in group-level COMMIT (ρ ≈ -0.8). To remove this strong obscuring effect, we recomputed correlations using residual edge weights following linear regression of edge length (**Figure 3B**). NoS, SIFT2 and COMMIT remain positively associated (group-level ρ ≈ 0.35) and R_1_ remains negatively associated with FC (group-level ρ ≈ -0.22). Correlation magnitude was reduced following linear regression in all cases. ICVF, FA and RD are reduced to 0 suggesting that they may not be useful in modeling whole-brain FC. These results support the idea that R_1_-weighted SC networks provide complementary information to NoS, SIFT2 and COMMIT about the brain structure-function relationship.

**Figure 3.**
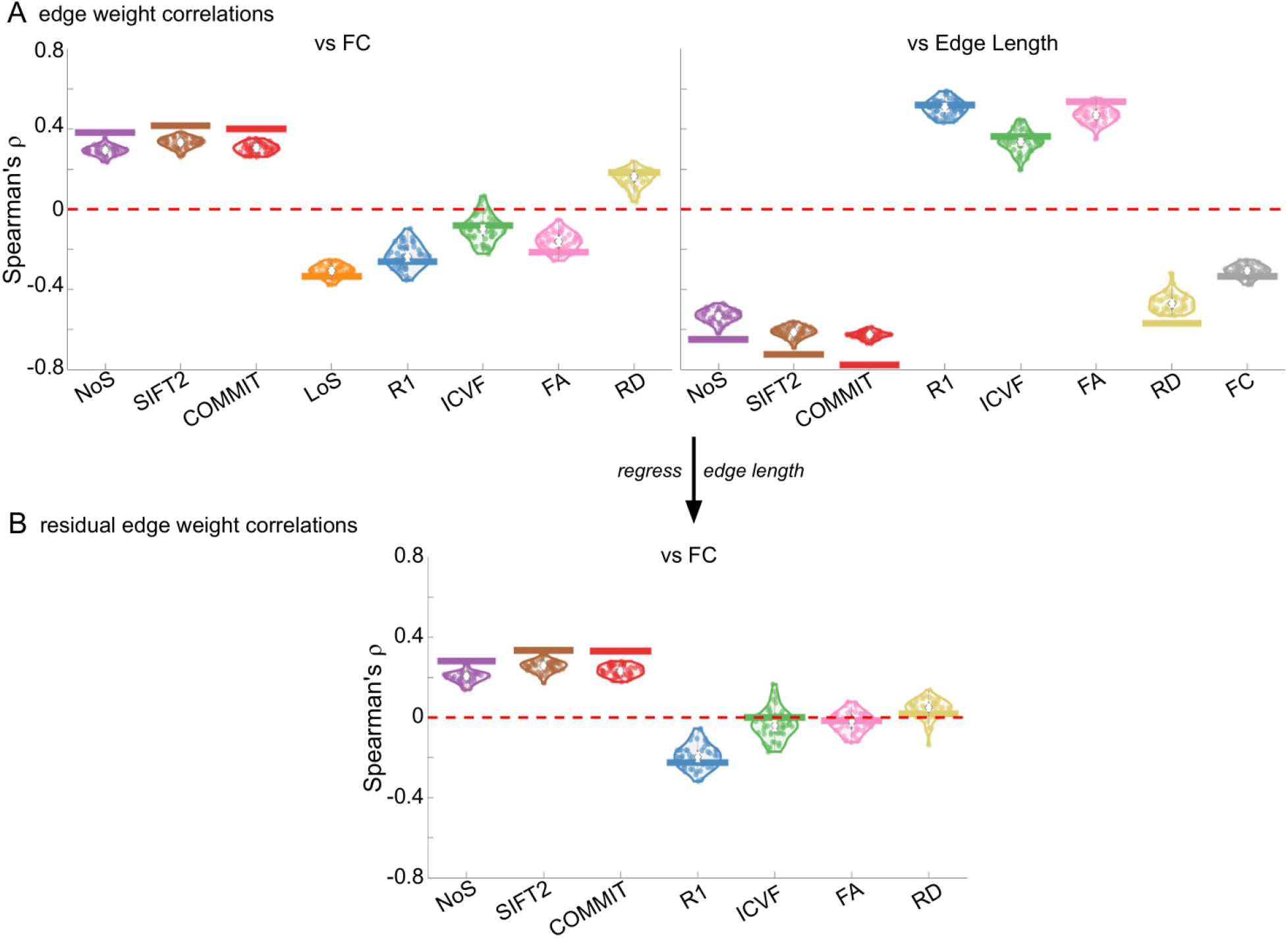
Edge Weight Correlations with FC and Edge Length. (A) Violin distributions of edgewise Spearman’s rank correlations of all networks with FC (left) and edge length (right). (B) Violin distributions of edgewise Spearman’s rank correlations of residual edge weights in all networks with residual edge weights in FC. Residual edge weights were computed by linear regression of edge length. Colored data points and bars respectively indicate subject-level and group-level correlations.

### Edge Caliber and Myelin Content are Inversely Related

Here, we ask how R_1_, which we refer to as the myelin-weighted network, is related to the connection-strength-weighted network COMMIT. Edge-length regressed residual edge weights in NoS, SIFT2 and COMMIT show a negative association with R_1_ residuals for all subjects and at the group level, which is strongest in COMMIT (group-level ρ & r ≈ -0.29) (**Figure 4A**). This suggests an edge-length independent inverse relationship between white matter structural features related to connection strength and myelin content.

**Figure 4.**
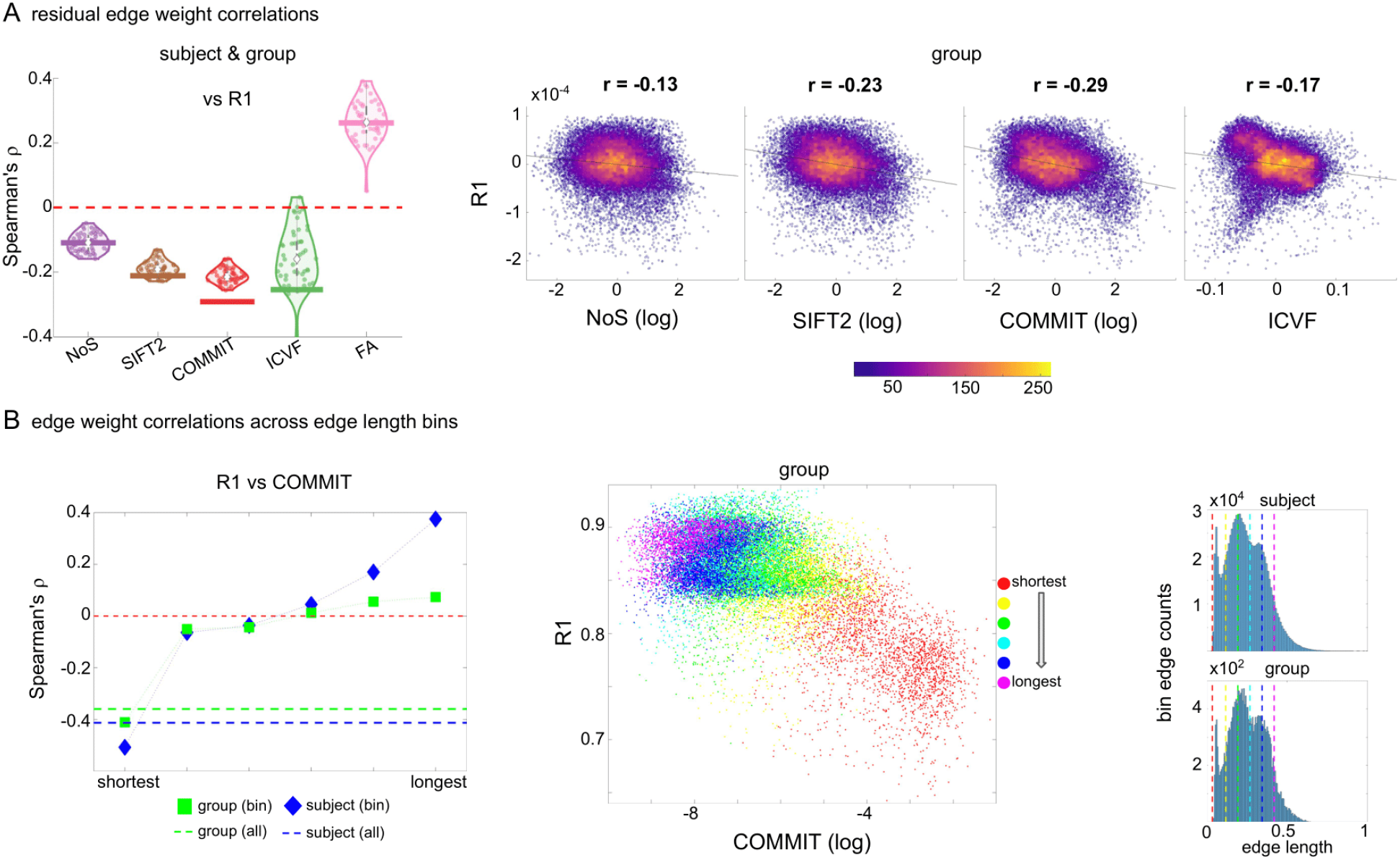
The Myelin-Dependence of Structural Brain Networks. (A) Violin distributions (left) of edgewise Spearman’s rank correlations with the myelin-weighted network R_1_. Residual edge weights are compared following linear regression of edge length. Colored data points and bars respectively indicate subject-level and group-level correlations. Heat scatter plots (right) of group-level residual edge weights in R_1_ as a function of NoS (left), SIFT2 (left middle), COMMIT (right middle) and ICVF (right) with the best fit linear curve shown in black. Color indicates data density. (B) Line plot (left) of edgewise Spearman’s rank correlation of edge weights in R_1_ vs COMMIT across edge length bins. Group-level and subject-level are respectively shown in green and blue. The square and diamond markers connected by dotted lines show binned correlation values, and the horizontal dashed green and blue lines mark the correlation values for all edges pooled together. Scatter plot (middle) of group-level edge weights in R_1_ as a function of COMMIT with data points colored by bin identity. Histograms (right) illustrating subject- and group-level edge length bins.

Computing correlations of edge weights (not residuals) within edge-length bins allows the inverse relationship between R_1_ and COMMIT to be traced to the shortest edges of the network (group ρ ≈ -0.40, subject ρ ≈ -0.50). As edge length increases, this relationship is reduced to 0, then becomes strongly positive in the longest subject-level edges (ρ ≈ 0.39). The scatter plot of group-level R_1_ vs COMMIT (middle) shows decreasing COMMIT and increasing R_1_ with increasing edge length. All together, these results support an inverse relationship between the edge caliber and myelin content of a given white matter tract. This can be partly explained by the differential dependence of these structural features on edge length: longer tracts tend to be more myelinated with lower total intra-axonal cross-sectional area. However, this relationship is robust to controlling for edge length supporting an intrinsic dependence between these white matter features.

### Divergent Small-Worldness, Hubness and Rich Club in Weighted Structural Networks

In this final section, we apply network analysis tools (Rubinov & Sporns, 2010) based on graph theory (Fornito et al., 2013; Sporns, 2018) to group-level weighted SC networks. This facilitates high-level interpretation of general features of network communication such as integrative vs segregative processing and the economy of network organization. Although the high material and metabolic cost of brain tissue naturally tends to favor local connectivity (high clustering), short overall network path length is achieved through a small number of relatively expensive long-range connections (Bullmore & Sporns, 2012). These edges and the nodes they interlink form a densely connected network core known as the rich club (Martijn P. van den Heuvel & Sporns, 2011). While the general proclivity for high local clustering gives rise to segregated functional modules, the rich-club nodes act as network communication hubs supporting inter-modular integration (Collin et al., 2014; de Reus & van den Heuvel, 2014; Griffa & Van den Heuvel, 2018; Kim & Min, 2020; Martijn P. van den Heuvel & Sporns, 2013). Thus, small-world network topology (high clustering and low path length) (Bassett & Bullmore, 2006, 2017) supports both integrative and segregative processing at a minimum of wiring cost, and the underlying scaffold of hub brain regions tend to show high centrality, low path length (high closeness) and low clustering (M. P. van den Heuvel et al., 2010).

Here, we report normalized small-worldness, normalized rich-club curves and nodal hubness (**Figure 5**). Normalized small-worldness (S) is computed as the quotient of normalized measures of clustering coefficient (C/C_null_) and path length (L/L_null_).

**Figure 5.**
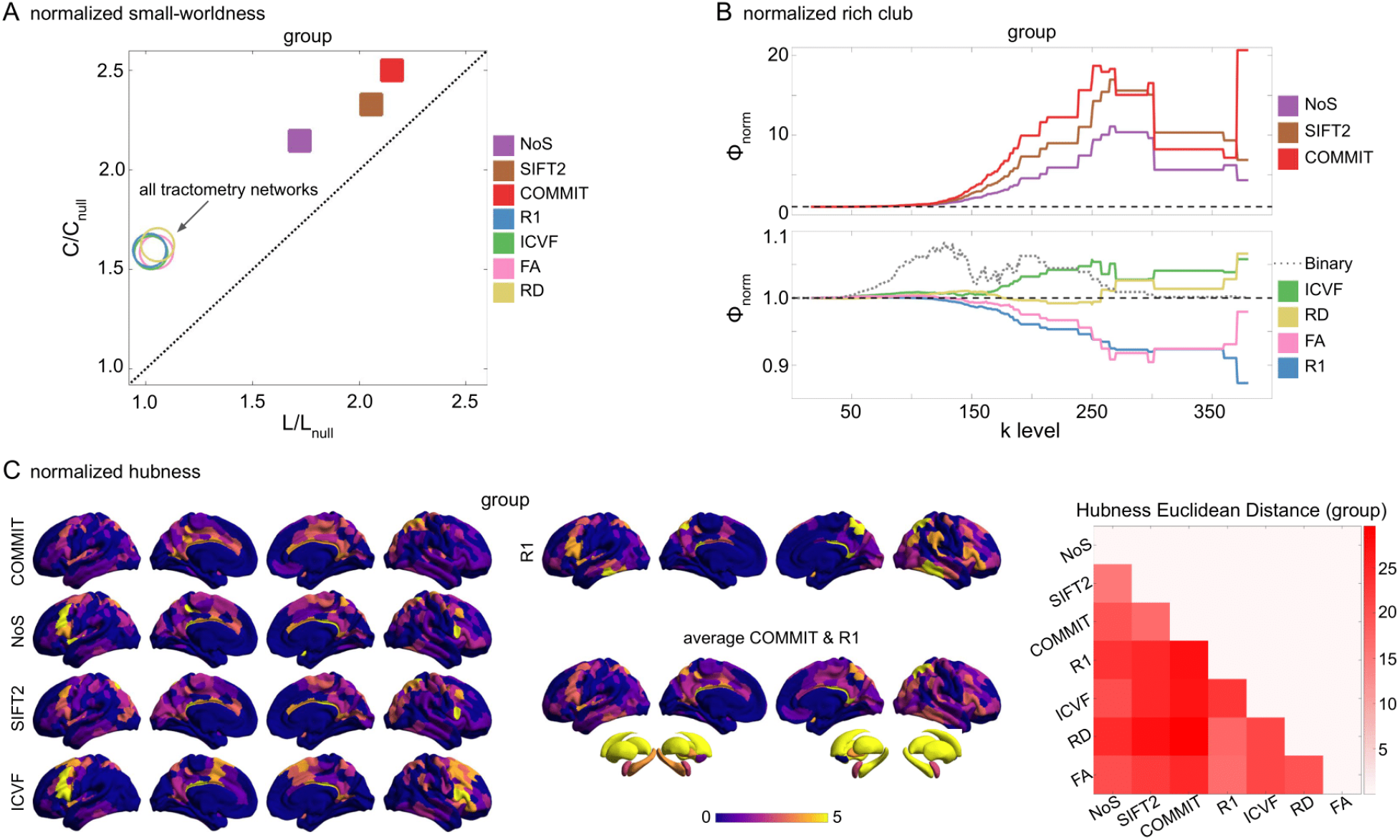
Group-level network topology. (A) Small-worldness was estimated in all structural networks: clustering coefficient was normalized within each node, averaged across nodes (C/C_null_), then plot as a function of normalized Characteristic path length (L/L_null_). Topology measures averaged across 50 degree and strength preserving null networks were used for normalization. Networks above the identity line (dotted black) are characterized by the small world attribute. Tractometry networks are indicated by the arrow. (B) Normalized rich club curves are shown for COMMIT, NoS and SIFT2 (top), as well as ICVF, RD, FA and R_1_ (bottom). A single binary network (dotted gray line) is also shown (bottom) as binary connectivity was uniform across weighted networks. The normalized rich club coefficient (ϕ_norm_) was computed across the range of degree (k) and normalized against 1000 null models (degree preserving for binary and degree and strength preserving for weighted networks). A ϕ_norm_ value > 1 (horizontal dashed black lines) over a range of k indicates the presence of a rich club. (C) Nodewise hubness scores are projected onto Schaefer-400 cortical and 14-ROI subcortical surfaces. Scores (0-5) were computed for each node as +1 point for all nodes in top 20% strength, betweenness, closeness and eigenvector centrality, as well as bottom 20% clustering coefficient. The matrix (right) shows the Euclidean distance between all pairs of nodal hubness vectors.

All group-level weighted SC networks show the normalized small-world property (S > 1) of higher clustering and lower path length than would be expected by chance (**Figure 5A**). Small-worldness is highest in COMMIT (S ≈ 2.5) and lowest in R_1_, ICVF, FA and RD (S ≈ 1.6). In contrast, all weighted SC networks did not show a canonical rich club (**Figure 5B**). Relative to the tractometry and binary SC networks, the normalized rich-club coefficient (ϕ_norm_) was much higher in magnitude in NoS, SIFT2 and COMMIT. A rich club was detected in these networks across a large range of degree (k) levels (150 < k < 300). ϕ_norm_ was maximal at k ≈ 265 in COMMIT. A rich club was also detected across a similar range of k levels in ICVF and across k in the range [250 300] for RD, albeit with much lower magnitude ϕ_norm_. However, no clear rich club was observed in R_1_ or FA. In fact, the rich-club curves for these networks are roughly symmetric about the ϕ_norm_ = 1 line relative to COMMIT. A densely connected core was of course recovered in all weighted SC networks (uniform binary connectivity), but these results suggest that its interconnecting edges were consistently weaker than would be expected by chance in R_1_ and FA. By comparison, a rich club was observed in the binary SC network across the very large range of k [50 300]. This supports two important concepts: (1) SC network edge weights can provide an additional layer of information useful for refining the topology of binary SC; and (2) different methods for computing SC network edge weights yield diverse network topology.

Weighted SC networks show network-dependent spatial topology of hubness scores (**Figure 5C**). The COMMIT and R1 averaged surface shows prominent hubs distributed throughout the brain including the fronto-parietal network. Nearly all of the subcortex showed a hubness score of 4 or greater in all networks. The Euclidean distance between hubness score vectors (right) was lower for COMMIT and SIFT2 than for either network with NoS. Of the streamline-specific networks, NoS was more similar to both R1 and IVCF. Overall, these results illustrate the considerable impact that edge weighting can have on network topology.

## DISCUSSION

Computational network modeling provides a customizable platform for probing the mechanistic relationship between human brain structure and function *in vivo*. Here, we assemble a thorough characterization of structural brain networks weighted by a range of quantitative MRI metrics capturing the macro- and microscopic features of white matter tracts to inform their utilization in computational models of brain function. Notable trends included: (1) greater edge weight contrast and skewed (heavy-tailed) distributions in the streamline-specific networks NoS, SIFT2 and COMMIT; (2) whole-brain correlations with FC in networks weighted by connection strength (positive) and myelin (negative) which were robust to controlling for edge length; (3) whole-brain inverse relationships with myelin for networks weighted by connection strength and neurite density independent of edge length; and (4) the absence of a rich club in R_1_ and FA networks. All weighted SC networks showed a strong spatial dependence and small-world architecture. Collectively, these results support the overall conclusion that SC networks weighted by edge caliber (e.g., SIFT2 and COMMIT) and myelin (e.g., R_1_) can be used to quantify non-overlapping subsets of white matter structural features related to FC supporting their joint utilization in modeling function.

### COMMIT vs SIFT2: The Superior Estimate of Connectivity Strength?

A principal goal of this work is to identify what, if any, advantage over NoS is provided by the global optimization methods SIFT2 and COMMIT. NoS has previously been used to inform the strength of interregional coupling in computational models of function (e.g., (Honey et al., 2009)). However, important limitations restrict model interpretation. Besides suffering from a range of biases related to the position, size, shape and length of white matter tracts (Girard et al., 2014), NoS varies as a function of tracking parameters limiting its specificity for white matter structural features (Jones, 2010; Jones et al., 2013). SIFT2 and COMMIT reportedly restore the quantitative link between connectome edge weights and white matter structural features related to connection strength. Our results show that when network density is uniform across structural metrics, COMMIT shows greater subject-specificity, edge weight contrast, correlation with myelin, small-worldness and rich club coefficient relative to SIFT2. This supports the hypothesis that using COMMIT instead of NoS to modulate the strength of interregional coupling in computational models of function will yield the greatest improvement in model fit.

### Myelin Complements Connection Strength in Predicting FC

Despite the differences between COMMIT, SIFT2 and NoS; our results indicate that their edge weights show roughly equivalent positive correlations with FC over the whole brain. R_1_ was negatively correlated with FC. Significant evidence indicates a link between cerebral myelin and FC including: a relationship between intracortical myelin and FC (Huntenburg et al., 2017; Wang et al., 2019); the prediction of cognition (Sonya Bells et al., 2017; Caeyenberghs et al., 2016) and FC-derived components (Messaritaki et al., 2021) with myelin-sensitive metrics; and a relationship between damaged myelin sheaths and greater conduction delays in multiple sclerosis (Sorrentino et al., 2022). At the cellular-level, myelin contributes to conduction velocity (Huxley & Stämpfli, 1949), metabolic support (Nave & Werner, 2014) and plasticity (Gibson et al., 2018), all of which could be argued to support brain function. Myelin plasticity in particular can be described in terms of “activity-dependence”, whereby an increase in the functional activity of a given circuit stimulates cellular signaling cascades promoting greater myelination (Douglas Fields, 2015; Mount & Monje, 2017). Coupled with our results, this complex mix of functional roles supports the idea that structure-function models will be improved by integrating measures of myelin and connection strength.

### Tract g-ratio and Edge Caliber are Inversely Related to Length

When controlling for edge length, we found an inverse relationship between R_1_ and COMMIT over the whole brain in all subjects and at the group level. This suggests that the aggregate g-ratio of a white matter tract may increase with edge caliber. At the cellular-level, the diameter of an axon and the thickness of its myelin sheath show nearly a linear relationship over a broad range of smaller diameter axons which becomes increasingly nonlinear as axon diameter increases (Berthold et al., 1983; Hildebrand & Hahn, 1978). In general, increasing axon diameter tends to outpace increasing myelin thickness i.e., g-ratio tends to increase with increasing axon caliber (Hildebrand & Hahn, 1978). Our findings suggest that this cellular-level principle may extend to the systems level: increases in edge caliber tend to outpace changes in the myelin content resulting in a concomitant increase in the g-ratio of white matter tracts.

We localized the inverse relationship between R_1_ and COMMIT to the shortest edges i.e., the g-ratio was the highest in the shortest connections. This result is supported by a previous imaging study showing the highest g-ratio in “local” connections (Mancini et al., 2018). In general, we found that R_1_ increased and COMMIT decreased with increasing edge length. Both of these trends fit well with theories of brain wiring economy in which the energetic cost of maintaining biological material increases with connection length (Bullmore & Sporns, 2012). This natural pressure acts to reduce the total axonal volume of longer white matter bundles. Increasing the myelin content of longer tracts comes at a cost as well, but this may be at least partially offset as increasing myelin content reduces the total membrane surface area along which expensive electrochemical gradients must be maintained (Bullmore & Sporns, 2012). Although, a cost-benefit analysis of the energetics of myelination concluded that the energetic cost of myelin maintenance outweighs any savings on action potentials (Harris & Attwell, 2012). This suggests that higher myelination of longer edges may be better explained as a mechanism to provide trophic support (Nave & Werner, 2014) to vital inter-regional connections (Martijn P. Van Den Heuvel et al., 2012) or to reduce conduction delays.

### Edge Weight Variance Decreases with Edge Length in Most Weighted Structural Networks?

White matter features related to myelin content, connection strength and neurite density tend to become more consistent across tracts as tract length increases. Greater variability in the weights of the shortest connections could result from a higher proportion of false positive streamlines influencing these edge weights. For SIFT2 and COMMIT, streamline weight computation becomes increasingly unstable with decreasing length as fewer voxels contribute to the fit. However, this result could also be explained more generally by contrasting the roles of shorter and longer connections in the brain. Shorter white matter tracts connect brain regions near each other in space e.g., within the same module. Just as we might expect the characteristics of smaller roads and streets (e.g., width, building materials, markings, signs, sidewalks, etc.) to vary by neighborhood and city, we might also expect the morphology of shorter white matter connections to change as the functional specialization of any given region or module changes. On the other hand, longer tracts (i.e., the freeways of the brain) may overlap more in both their functional role and morphological features relative to shorter connections, hence lower edge weight variability. Breaking with the above pattern, FA and RD showed the highest edge weight variance in the longest connections. Given that structural measures derived using a voxel-wise diffusion tensor model are particularly sensitive to the white matter “architectural paradigm” (Jones et al., 2013), these results suggest that white matter features related to fiber orientation and geometry actually diverge with increasing tract length.

### The Absence of a Rich Club in R_1_ and FA

Group-level R_1_ and FA did not show a normalized weighted rich club for any degree k. Higher myelination in the white matter tracts connecting rich club nodes has previously been reported (Collin et al., 2014); however, methodological differences limit comparability. A rich club has previously been reported in FA-weighted networks using similar methods to ours (Martijn P. van den Heuvel & Sporns, 2011). The source of this disagreement could potentially be attributed to differences in our tractography algorithm, parcellation or null network computation.

In weighted rich-club detection, the identification of a densely connected core is independent of edge weight (depends only on node degree), but the designation of this subnetwork as a rich club requires that it contains a higher-than-chance proportion of the strongest edges from the full network. Indeed, this is the case over a broad range of degree k for COMMIT. Over the same range of k, the normalized rich-club curves for R_1_ and FA are inverted about the threshold value of 1 with respect to COMMIT. This implies that the subnetwork found at a given k in this range contains edges which tend to show higher COMMIT and lower R_1_ edge weights than expected by chance. We previously showed edgewise inverse correlations between R_1_ and COMMIT which were robust to controlling for edge length. We also showed that R_1_ and FA are positively correlated under these same conditions. In this light, it is not surprising that the edges connecting rich-club nodes tend to show opposite trends in R_1_- and FA-weighting with respect to COMMIT. Nonetheless, it is possible that the lack of a rich club in our myelin-weighted network is an artifact of tractometry. Future work will attempt to replicate this result using myelin-weighted networks computed with a different methodology (Schiavi et al., 2022).

### Limitations

Streamline tractography is known to suffer from several important biases including both false positive and negative streamlines which can influence downstream analyses (Maier-Hein et al., 2017; Schilling et al., 2019; Sotiropoulos & Zalesky, 2019; Zalesky et al., 2016). Through probabilistic tracking, we opted to minimize false negatives while maximizing false positives. This allowed us to implement careful streamline- and edge-filtering strategies in post-processing to address this known bias. Still, without a ground truth, we cannot quantify the extent to which we were successful in mitigating this issue, nor can we guarantee that we did not erroneously filter true positive streamlines or edges. All processing and filtering methods were consistent and network density was uniform across weighted structural networks. Thus, any major tractography bias should be as homogeneous as possible across networks.

Tractometry-derived brain networks suffer from widespread partial volume effects. The net effect of this bias is well understood and is apparent in our results and previous work (De Santis et al., 2014; Schiavi et al., 2022). Nonetheless, this method was included here as our goal was to characterize widely used structural connectivity methods. New techniques for reducing this bias are currently being developed which allow for the estimation of tract-specific microstructural features (e.g., (Barakovic, Girard, et al., 2021; Barakovic, Tax, et al., 2021; De Santis et al., 2016; Leppert et al., 2021; Schiavi et al., 2022)).

### Conclusion

We presented a thorough characterization of weighted SC networks. Overall, our findings support the joint use of SC networks weighted by connection strength and myelin in predicting FC. In particular, using the COMMIT algorithm to quantify connection strength shows promise. Beyond R_1_, there are a wide array of myelin sensitive metrics that could be used to compute useful myelin-weighted networks. The integration of this microstructure-weighted connectivity approach into structure-function models will advance the mechanistic interpretation of both the function and dysfunction of the living human brain.

## MATERIALS and METHODS

These data are available for download (https://portal.conp.ca/dataset?id=projects/mica-mics). See Royer et al. (Royer et al., 2022), Cruces et al. (Cruces et al., 2022) for full details of data acquisition and processing. All data processing and analysis code is openly available at https://github.com/TardifLab/Weighted-SC-Networks.

### Data Acquisition & Preprocessing

Multimodal MRI data was collected in 50 healthy volunteers at 3 Tesla as follows:

- T_1_-weighted (T_1_w) anatomical: 3D magnetization-prepared rapid gradient-echo sequence (MP-RAGE; 0.8mm isotropic)
- Multi-shell diffusion-weighted imaging (DWI): 2D pulsed gradient spin-echo echo-planar imaging sequence (1.6mm isotropic); three shells with b-values 300, 700, and 2000s/mm^2^ and diffusion directions 10, 40, and 90
- 7 minutes of resting-state functional MRI: multi-band accelerated 2D-BOLD gradient echo echo-planar sequence (3mm isotropic)
- A quantitative T_1_ map: 3D-MP2RAGE sequence (Marques et al., 2010) (0.8mm isotropic)

The multi-modal processing pipeline *micapipe* (Cruces et al., 2022) (https://micapipe.readthedocs.io/) was used to preprocess diffusion, anatomical, and functional images. Functional data derivatives were obtained in parcellated FC matrix form.

### Tractography and Microstructural Metrics

To estimate structural connectomes, anatomically constrained tractography (R. E. Smith et al., 2012) was performed on the normalized white matter FOD image using the probabilistic algorithm iFOD2 (J.-D. Tournier et al., 2010). Tractograms of 5 million streamlines were generated by seeding the gray-white matter interface using the following parameters: maxlength=400, minlength=10, angle=22.5, step=0.5, cutoff=0.06, backtrack, crop_at_gmwmi (gray-matter-white-matter interface). These tractograms were filtered in a two-stage process. (1) a whole-brain connectome weighted by NoS was computed then decomposed into its composite streamlines to derive a new tractogram in which any streamline which failed to connect two gray matter ROIs was excluded. This “streamline-filtering” step typically resulted in approximately a 5% decrease in the size of the tractogram (∼250k streamlines removed) and was undertaken to ensure that these erroneous streamlines did not affect the COMMIT model. Streamline-filtered tractograms were used to compute NoS and were used as inputs to both the SIFT2 and COMMIT models. SIFT2 determines the effective cross-sectional area of each streamline such that the streamline density throughout the white matter fits the fiber densities estimated using spherical deconvolution. COMMIT was run using a Stick-Zeppelin-Ball forward model and default settings (see https://github.com/daducci/COMMIT). Using the simplifying assumption that structural features are constant along the length of a streamline, COMMIT can be used to compute a weight for each streamline representing their respective proportion of the global diffusion signal i.e., the cross-sectional area of their intracellular compartment. (2) Any streamline with a COMMIT weight < 1e^-12^ (machine precision 0) was interpreted as a false positive and filtered from the tractogram. This streamline-level COMMIT-filtering step typically resulted in greater than a 90% decrease in the size of the tractogram with most containing between ∼300-600k streamlines. COMMIT-filtered tractograms were used not only in the computation of COMMIT, but all tractometry networks as well. This additional filtering step was performed on COMMIT streamline weights only (not SIFT2) to reduce the impact of false positive streamlines in tractometry networks as much as possible.

### Construction of Weighted Structural Networks

The streamline-specific SC networks were computed in the following manner: (1) NoS as the summed streamline count; (2) LoS as the mean streamline length; (3) SIFT2 as the sum of SIFT2 streamline weights; and (4) COMMIT as the length-weighted sum of COMMIT streamline weights as in (Schiavi et al., 2020). Explicitly, edgewise entries in COMMIT-weighted networks were computed as:

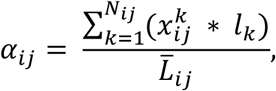

where *α*_*ij*_ is the edge weight between nodes *i* and *j*; 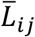 is the mean streamline length; *N*_*ij*_ is the number of streamlines; 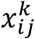 is the COMMIT weight of streamline k; and *l*_*k*_ is its length. Edge weights in NoS, SIFT2 and COMMIT were normalized by node volume.

SC networks weighted by FA, RD, ICVF (Zhang et al., 2012) and R_1_ were derived using multi-modal tractometry (S Bells et al., 2011). Streamline weights were computed by: (1) co-registering the tractogram and desired image; and (2) sampling the voxel-level aggregate value along the length of each streamline. Edge weights were computed as the median along each streamline and the mean across streamlines by node pair. Voxel-wise measures of FA and RD were computed with a diffusion tensor model (Basser et al., 1994) and ICVF by applying the NODDI multi-compartment model (Zhang et al., 2012) to preprocessed DWI data (Daducci, Canales-Rodríguez, et al., 2015).

The 400-node Schaefer (Schaefer et al., 2018) cortical parcellation is used in all results. Subcortical ROIs corresponded to 7 bilateral regions (14 nodes) including the amygdala, thalamus, caudate, accumbens, putamen, hippocampus, and pallidum. A single static, zero-lag FC network was derived by product-moment pairwise Pearson cross-correlation of node-averaged time series. FC network edge weights were Fisher Z-transformed.

### Connectome post-processing

All SC networks were thresholded at the edge level within subject by: (1) setting edges = 0 in all weighted SC networks if they had a COMMIT edge weight < 1e^-12^; and (2) applying a 50% uniform threshold mask to facilitate group-consensus averaging. This minimized differences in binary structural network density across subjects and enforced uniform density across weighted SC networks at the group level and within subject. COMMIT was used for this filter as it had the lowest connection density to start.

### Network Analysis

Network analysis was performed using tools (Rubinov & Sporns, 2010) based on graph theory (Fornito et al., 2013; Sporns, 2018). Measures of clustering coefficient and path length were normalized against 50 degree and strength preserving null networks. Clustering coefficient was normalized within node then averaged across nodes to obtain a scalar value per network. The following weight (W_ij_) to length (L_ij_) transform was used in path length computation: L_ij_ = - log(W_ij_). Weighted rich club curves were normalized against 1000 degree and strength preserving null networks. The edges in all degree and strength preserving null networks were rewired 1e^6^ times total, and the strength sequence was approximated using simulated annealing. Rich club curves were normalized in binary networks against 1000 degree preserving null networks in which each edge was rewired 100 times. All edge rewiring followed the Maslov & Sneppen rewiring model (Maslov & Sneppen, 2002). Similar to (M. P. van den Heuvel et al., 2010), hubness scores (0-5) were computed as 1 point for all nodes showing top 20% strength, betweenness, closeness or eigenvector centrality; and lowest 20% clustering coefficient.

## ACKNOWLEDGMENTS

We acknowledge research support from the National Science and Engineering Research Council of Canada (NSERC Discovery-1304413, DGECR-2018-00216), the CIHR (FDN-154298, PJT-174995), SickKids Foundation (NI17-039), Azrieli Center for Autism Research (ACAR-TACC), Brain Canada, Fonds de recherche du Québec – Santé (FRQ-S), Healthy Brains for Healthy Lives, and the Tier-2 Canada Research Chairs program.

## Notes

https://github.com/TardifLab/Weighted-SC-Networks

https://portal.conp.ca/dataset?id=projects/mica-mics

### Competing Interest Statement

The authors have declared no competing interest.

